# Pericytes in heart

**DOI:** 10.1101/2025.05.14.654087

**Authors:** Guiling Zhao, W. Jonathan Lederer

## Abstract

Pericytes are cells associated primarily with capillaries and are thought to play an important role in the regulation of blood flow. They are often referred to as “mural” cells because they are so frequently found on the exterior walls of small vessels - particularly the capillaries. In heart, high-resolution real-time observations and measurements of pericyte function under physiological conditions are challenging to obtain because of vascular motion, tissue depth and vigorous functional movement. For these reasons, the heart may be one of the most difficult tissues in which to examine pericyte function. Recently, we introduced a perfused papillary muscle preparation (the Z-Prep) that allows us to observe coronary arteries, arterioles, venules, capillaries and myocytes in real time at physiological temperature and pressure while also imaging pericytes ^1, 2^. Here we present an initial study intended to visualize and characterize quantitatively cardiac pericytes in heart at physiological pressure and temperature conditions. Vascular anatomy was imaged using a z-stack protocol with a rapidly spinning disk confocal microscope. Here the anatomical organization of the pericytes is shown at high resolution with respect to the microcirculation components and cardiac myocytes. The surprising findings include the high abundance of pericytes in native tissue, the extent of their spread on the capillaries themselves, and the existence of major pericyte extensions that travel intimately along the surface of neighboring ventricular myocytes and attach to capillaries on the distant side. These extensions arise from a capillary-based pericyte location and normally end on another capillary endothelial surface and we have named them “bridging” pericytes. Taken together this anatomical organization suggests that the pericytes provide signaling, communication and contractile services to important cellular components of the heart. There is also a provocative suggestion that pericytes in heart are unusually fragile since they suffer an extremely high degree of loss during cellular isolation procedures. However, our investigation of the organization argues against this fragility because of the durability of the dynamic pericyte organization and function despite the stress and brutality of the contracting heart. The work presented here lays the foundation for critical functional studies of pericytes in heart in both health and disease.

## Introduction

Studies of cardiac capillary dynamics and pericyte function present a formidable challenge in cardiovascular research. Within the complex landscape of the microvasculature in heart, understanding the capillary functional organization is of utmost importance. Here we attempt to extend our understanding of the roles played by pericytes. It has been hypothesized, for example, that the rapid and accurate control of local blood flow by small vessels in the heart is critical for healthy normal heart function, that it is driven by the metabolic state of the contractile tissue and that the capillaries themselves communicate this signal to contractile elements that include the pericytes and the vascular smooth muscle cells on arterioles. This process has been dubbed “electro-metabolic signaling (EMS)” ^2-4^. However, many aspects of EMS remain unclear due to the inherent complexities of studying these components in the beating heart. The relentless motion and rhythmic contractions of the heart make in vivo and ex vivo investigations exceptionally demanding (see reviews ^5-7^). Furthermore, the pursuit of real-time imaging and observation of these structures within the dynamic cardiac environment pushes the boundaries of current imaging technological capabilities. Consequently, most of our current knowledge about coronary microvessels and their pericytes comes from fixed, static heart tissues ^8-11^. A lot of elegant very recent work has already revealed the sophisticated structures of the heart microcirculation and the local architecture of the mural cells (e.g. pericytes, vascular smooth muscle cells). Such recent work has advanced our knowledge of the cardiovascular microcirculation biology ^11-13^. However, the process of tissue fixation can cause damage, deformation, and loss of cells and other components, leading to possible inaccurate observations and data interpretation. Additionally, one-time snapshot images cannot capture the dynamic changes that occur under diverse conditions including those that are stressful and challenging. Certainly, our understanding based on past studies of physiological and pathological changes in the microcirculation (based primarily on fixed tissue) should be verified. Thus, efforts to image live tissues in vivo and ex vivo have also been ongoing and are important ^5, 14-18^. Significant advances in live-tissue imaging have been achieved in brain research in recent years ^19-21^. However, less progress has been made in heart research due to the non-stop beating of the heart and the difficulty of performing surgery to allow long-term imaging of the dynamics of its vasculature.

To investigate active structural and functional changes in the heart, we recently developed a mouse papillary muscle preparation (Z-Prep) that allows us to pressurize the blood vessels and perfuse the tissue ^1, 2, 4^. This preparation enables high-resolution three-dimensional visualization of the microvascular bed. Using the Z-Prep, we successfully captured the microvascular diameter change in the presence of vasodilators and vasoconstrictors ^1, 2^. On the other hand, confusion remains regarding the identification of different mural cells and their respective functions in the heart. A more detailed characterization of the morphology of different mural cell types and the microvascular territories they occupy is needed to aid in the interpretation of the functional behavior of heart pericytes, both physiologically and pathologically. In this study, we aim to describe and advance our understanding of the structures of the “real” microvascular beds and illustrate the architecture of the surrounding components with an emphasis on pericytes. Using NG2-DsRed and PDGFRβ-tdTomato transgenic mice, we have characterized and categorized pericytes according to their locations, morphologies, and the presence of smooth muscle actin. Although the classification of pericytes has historically been challenging due to their phenotypic plasticity and the overlap of marker expression with other cell types, our efforts have provided important clarity. Finally, the function of pericytes in the regulation of microvascular blood flow is discussed based on the results presented here.

## Methods

### Preparation of arterially perfused mouse right ventricle papillary muscle ^1, 2^

The pressurized and arterially perfused mouse right ventricle papillary muscle was prepared as described previously ^1, 2^. In brief, the heart of the wild type or transgenic mouse (NG2-DsRed and PDGFRβ-tdTomato) under anesthesia was removed and placed into ice-cold Tyrode’s solution (in mM: 140 NaCl, 10 HEPES, 0.5 MgCl_2_, 0.33 NaHPO_4_, 5.5 Glucose, 1.8 CaCl_2_, and 5 KCl (pH 7.4 with NaOH)) containing 30 mM BDM (2,3-Butanedione monoxime). After cleaning of the connective tissues, the heart was transferred into the pre-chilled chamber that has been coated with PDMS (polydimethylsiloxane) and filled with Tyrode’s solution containing BDM (30 mM). Then the right ventricle was cut open carefully and the right ventricular free wall was removed. The septal artery was exposed. A nylon thread (30 μm in diameter, Living Systems Instrumentation, Burlington, VT, USA) was placed under the sepal artery and a loose knot was tied. The left ventricular free wall was removed, and the sample was transferred into the PDMS-coated experimental chamber (filled with Tyrode’s solution containing BDM) in which the cannula was positioned. The septal artery was then cannulated, and the small pins were used to secure the papillary muscle to the chamber floor so that the microscope objective had a clear view of the vasculature. Then the cannula was connected to a pressure/flow control pump or a gravity column to allow pressurized perfusion into the vascular lumen. The bath was superfused with physiological saline solution (PSS, in mM: 112 NaCl, 5 KCl, 1.2 MgSO_4_, 1.2 NaH_2_PO_4_, 24 NaHCO_3_, 10 glucose, 1.8 CaCl_2_, bubbled with 5% CO_2_) at ∼2 to 3 ml/min at 35-37°C.

### Dye loading and imaging on Z-Prep

After ∼30 min stabilization, Alexa Fluor-488 or 633 conjugated wheat germ agglutinin (WGA, 20 μg/ml) or Isolectin B4 (IB4) was loaded into the arterial lumen perfusate for ∼30 min followed by a complete wash. The papillary muscle (or septum) was first located visually and imaged using a 4x objective. At that point a high-power objective (40x) was used to image capillaries and pericytes. A spinning disc or point scan confocal was used for high resolution imaging. For photo bleaching of DsRed, the 561 nm laser power was briefly increased to 100%. Biochemical (i.e. endothelin-1 [ET-1] and BDM) and dyes of choice were perfused through the Z-prep arterial cannula along with other components of the perfusate. The perfusion pressure was monitored using a pressure transducer as needed and as previously reported ^1, 2^. Unless specified otherwise, BDM (30 mM) was included in the perfusates to minimize the motion of the tissue.

### Whole Mount Staining and Immunostaining Procedure

NG2-DsRed mouse hearts were cannulated and perfused using the Langendorff perfusion system with Ca^2+^-free Tyrode’s solution at room temperature. IB4 (10 μg/mL), Mitotracker, and Hoechst stain (10 μM) were loaded for 30 minutes, followed by a 10-minute wash. Subsequently, 4% paraformaldehyde (in PBS) was perfused through the vasculature for 15 minutes to fix the heart, and this perfusion was followed by a 5-minute wash using PBS. The heart was then removed from the Langendorff perfusion rig, and imaging was performed on either the ventricles or the papillary muscle. For whole mount immunostaining, after dye loading, wild-type hearts were permeabilized using 0.1% or 0.2% Triton X-100 (in PBS) for 15 minutes. The heart was removed from the Langendorff perfusion rig, and the right ventricular papillary muscle was dissected for whole-mount immunostaining. The tissue was incubated with primary antibodies—anti-NG2 (MAB5384, 1:100; Millipore Sigma-Aldrich) and anti-SMC-actin (Ab5694, Abcam)—for over two nights (∼36 h) at 4°C in a blocking solution (5% goat serum and 0.1% or 0.2% Triton X-100 in PBS), followed by a 2-hour wash with PBS. The tissues were then incubated with an Alexa Fluor-conjugated secondary antibody (1:200 to 1:3,000) for 4 to 6 hours at room temperature, followed by an overnight wash with PBS at 4°C. The tissues were then either mounted on a Sylgard-coated chamber using a fine needle or glued onto a coverslip and imaged using upright confocal microscopy (Nikon A1R).

### Image processing and Data analysis

The images or data (3D image renderings, measurements of vessel diameter, capillary segment length, pericyte distance, etc.) were processed and analyzed using either Imaris or Nikon software (NIS) for general analysis, version 3 (GA3). For vessel diameter and capillary segment length measurements, we drew lines manually across the vessel width (for diameter) and along the capillary segment length using 3D-rendered vessels. The volumes of imaged tissue, capillaries and pericytes were measured by 3D thresholding using GA3 function in NIS. The cellular processes and the capillaries were processed to ensure all were captured in the thresholding images. The volume ratio was calculated over the imaged tissue volume. Similarly, the myocyte volume was measured on the ones that was fluorescently injected. Statistical analysis was conducted using Origin 2018, with a significance value of P<0.05.

## Results

### Perfused right ventricle papillary muscle preparation - Z-Preps

Almost all experiments on living tissue were performed using Z-Preps, as illustrated in **Figure 1**. This figure shows a perfused and pressurized mouse right ventricle papillary muscle preparation ^1, 2^. Typically, the septal artery originates from the right coronary artery (illustrated in **Figure 1A**). However, in some instances, it can arise from the left coronary artery ^22^. The arterially perfused septum and papillary muscles were studied at physiological pressure, ranging from 40 mmHg to 80 mmHg. Room temperature or physiological temperature (∼35 to 37 °C, functional investigations) were applied through an inline heater. Fluorophore-conjugated lectins, such as wheat germ agglutinin (WGA) or IB4, were utilized to visualize the vessel beds. At lower magnifications (using a 4x objective lens), two or more papillary muscles within the same field of view could be observed, as shown in **Figure 1B** and **1C** ^1, 2^. These papillary muscles exhibited varying shapes: the first one appeared relatively round; the second elongated, and the third was also elongated but smaller (**Figure 1B** to **1D**). **Figure 1C** presents a typical Z-Prep, showcasing the septum and mouse papillary muscles. Due to its proximity to the cannula, the first papillary muscle is a feasible choice for imaging and data collection. However, the second papillary muscle is often preferred (**Figure 1B** to **1D**) because it is comparatively thinner, smaller, flatter, and better perfused. These characteristics facilitate more straightforward and consistent imaging (**Figure 1D**). The third papillary muscle which is located deeper was more challenging to keep in focus and thus less frequently studied. On occasion, well-perfused areas of the septum were also imaged.

**Figure 1.**
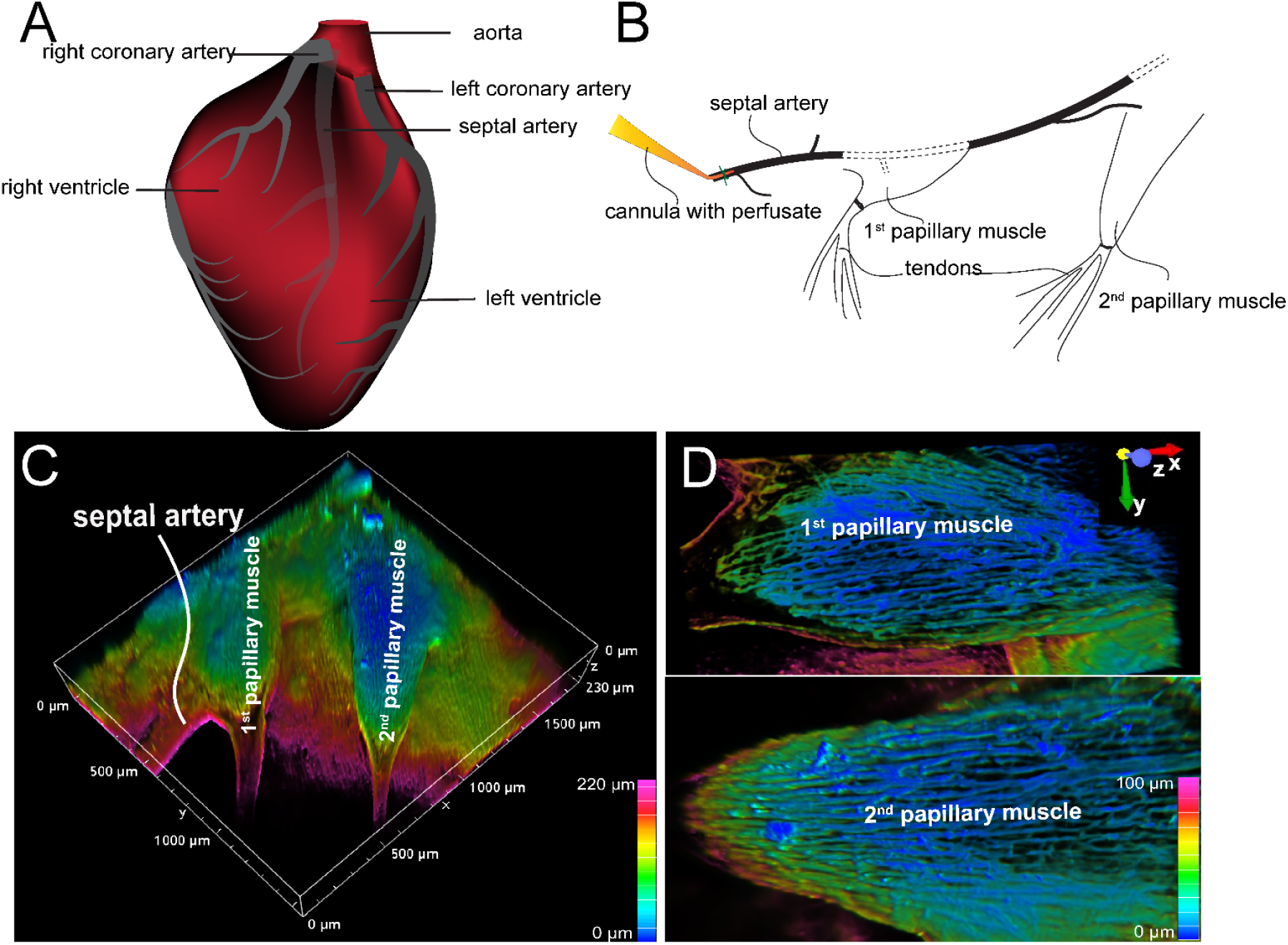
Arterially perfused mouse right ventricle papillary muscle. **A**. Cartoon shows the origin and course of coronary arteries. Note: septal artery originates from right coronary artery. **B**. Diagram shows the location of cannula and the papillary muscles. **C**. Depth-encoded vascular imaging of a Z-prep using WGA conjugated with Alexa Fluor 488 as fluorescent marker of the vasculature over a depth of 100 micros. **D**. Zoomed-in images from **C** with 1st (top) and 2nd papillary muscle (bottom). Depth encoding over 100 microns.

### Capillary network geometry

*The vascular organization*. We tracked the septal artery to examine the arterioles and capillaries. However, only the superficial arteries and arterioles and their offshoots were able to be examined (**Figure 2**) using monophoton confocal or spinning disc confocal imaging. The daughter branches were observed using Z-stack with 0.75 to 1 μm intervals. The capillary orders are not easy to track due to the depth and dense packed arrangement. Importantly, the organizational hierarchy of capillaries in heart differs significantly from that in the brain and retina ^4, 23, 24^, where capillaries are classified according to their order in a sequential branching pattern. Instead, the capillaries in the heart form a dense and simpler mesh-like network. This network is characterized by extensive branching and anastomoses, with longer capillary segments oriented parallel to the cardiac myocytes. Therefore, it is easy to identify and distinguish capillaries from precapillary arterioles, but it is difficult to distinguish between mother and daughter branches.

**Figure 2.**
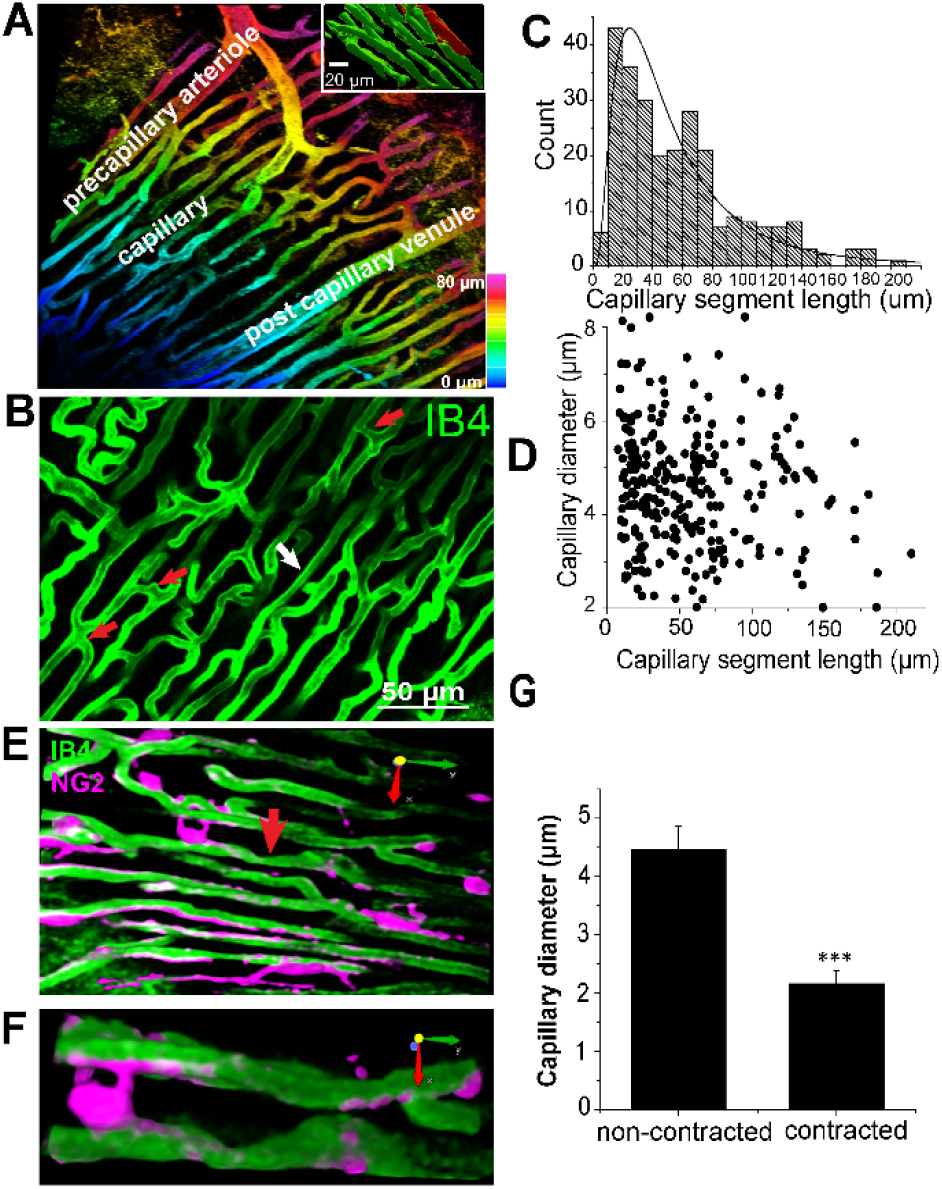
Capillary network geometry and the size in Mouse Heart (Z-Prep). **A**. A 3D image with colored depth coding illustrates the organizational hierarchy of precapillary arterioles, the capillary bed, and the postcapillary venule. The inset binary image shows the orientation of capillaries (green) parallel to the myocyte (red). The myocyte was filled with sulforhodamine through microinjection. **B**. The capillary bed displays varying lengths of capillary segments, with shorter segments indicated by red arrows (< 10 μm) and longer segments by a white arrow (>100 μm). **C**. A histogram depicts the distribution of different capillary segment lengths. **D**. A correlation scatterplot of capillary diameter and segment length indicates no significant relationship between the two. **E**. Examples demonstrate capillary local constriction (arrow) and the location of pericytes (magenta). **F**. A zoom-in of the arrow-pointed area in **E. G**. Summary data showing the capillary diameter of non-contracted and local contracted area.

### Capillaries

As shown in **Figures 2A** and **2B**, the orientation and size of the precapillary arterioles are distinctly different from those of the capillaries. Since almost all the capillaries align tightly and longitudinally with the cardiac myocytes immediately after their branching from the arterioles, we speculate that the diameters of all the capillaries are similar regardless of the segment length. Indeed, the capillary diameter measures 4.96±0.1 μm under perfusion pressures ranging from 40 to 60 mmHg in the presence of BDM (30 mM). This number is smaller than the corrected value from the earlier work on fixed tissues ^9, 13^. The averaged capillary segment length was measured to be 58.7±3.8 μm (**Figure 2C**), with the shortest segment measuring less than 10 μm, and the longest exceeding 180 μm (**Figure 2B** to **2D**), which is considerably shorter than what was observed in rat heart and skeletal muscle ^25, 26^, but similar to pig heart capillary ^13^. At the present time these length differences do not appear to be functionally important. **Figure 2D** demonstrates that the capillary segment length does not necessarily correlate with the diameter of the capillary. The volumes of the imaged tissue and the capillaries were also analyzed offline using binary thresholding and 3D length measurements provided by NIS software and Imaris. Using this analysis, the capillaries occupy approximately 13.5±0.8% of the total tissue volume.

### Capillary diameter dynamics

Activation of K_ATP_ channels in perfused heart tissue (Z-prep) leads to capillary dilation^2^ when we perfuse at 40-60 mm Hg at a physiological temperature. We showed this by adding the K_ATP_ opener thus induces capillary dilation^2^ and this produced an increase in blood flow. We deduced that this vasodilation was due to the K_ATP_-dependent electrical time-averaged hyperpolarization of ventricular myocytes and thus other elements in the vascular tree including the capillary endothelial cells, the pericytes and the vascular smooth muscle cells on the end arterioles^2^. Increased capillary perfusion pressure to 100 mm Hg in the absence of a vasodilator also increases blood flow and capillary diameter (from ∼4.0 to ∼4.5 microns). We also show that a K_ATP_ channel inhibitor produces a reduction in blood flow and in capillary diameter ^2^. Local constriction or narrowing is frequently observed in our preparation, especially in the absence of BDM (**Figure 2E** to **2G**). The diameter of the narrowest area in pressurized vascular systems is ∼ 2 μm (**Figure 2G**). The distance of the narrow center to the pericyte soma is 29.3±5.7 μm, suggesting that pericyte cell soma is not directly involved in all capillary local constrictions. The capillary diameter can be altered in pathologic situations such as ischemia ^10, 27^. Pathologically, capillary closure is observed in heart, brain and other tissues ^10, 28, 29^, which can decrease or even stop blood flow - when it occurs. Capillary dilation was also observed in the living tissues other than heart, especially in the presence of vasodilator or when the tissue was exposed in acute hypoxia ^27, 30-32^.

### Structural Diversity of Cardiac Pericytes

*Intimacy: pericytes and myocytes*. In this study, DsRed-tagged NG2 or tdTomato-tagged PDGFRβ transgenic mice were employed to identify pericytes (**Figure 2E-2F, 3A** to **3D, Figure 4A** and **4D, Figure 5A, Figure 6A**). Despite the expression of both NG2 and PDGFRβ in arterial smooth muscle cells (ASMCs), these markers proved sufficient for effectively distinguishing pericytes from ASMCs using Z-Prep ^1-4, 30, 33^. As depicted in **Figure 3** and supported by our prior research ^1, 2^, pericytes were readily distinguishable from ASMCs based on their distinct location within the capillary beds and their characteristic cellular morphology^1, 2, 4^. In the cardiac microenvironment, the compact and tight packing of myocytes results in shorter capillary segments between myocyte layers and many bifurcation angles closer to 90 degrees (**Figure 3Af**). Consequently, the spatial arrangement of pericyte somas and processes becomes particularly intriguing and provocative (**Figure 3E** and **3F**). Even though it is challenging to identify the location and spread of each pericyte with respect to the other cells that surround it, it can be done. We used a simplification to estimate 3 pericyte locations each with a distinct character. 1. 12.0±2.6% of the pericytes were identified as those located at the junctional areas where capillaries bifurcate (**Figure 4Aa, 4B**). 2. 12.2±3.0% of the pericytes were located on the capillaries themselves. 3. 75.8±4.5% of the pericytes formed a bridge between at least two capillaries. Examples of these three types are shown in **Figures 4A and 4B**. While all the pericytes are likely to have regions of exposure to ventricular myocytes, the “bridging pericytes” dominate in their exposure to and contact with ventricular myocytes! In addition, this is the predominant pericyte type. As the bridging pericytes stretch from one capillary to two or more capillaries (**Figure 4D** and **4E**), much of the cell body and the connecting arms are surrounded closely by the sarcolemmal membranes of the ventricular myocytes. In many cases there are regions of intimate juxtaposition between such pericytes and their nearby myocytes. This spatial organization of the cardiac pericytes is thus very different from the capillary-centric localization pattern of pericytes observed in the retina and other parts of the brain ^23, 31, 34^ and has not been reported before to the best of our knowledge.

**Figure 3.**
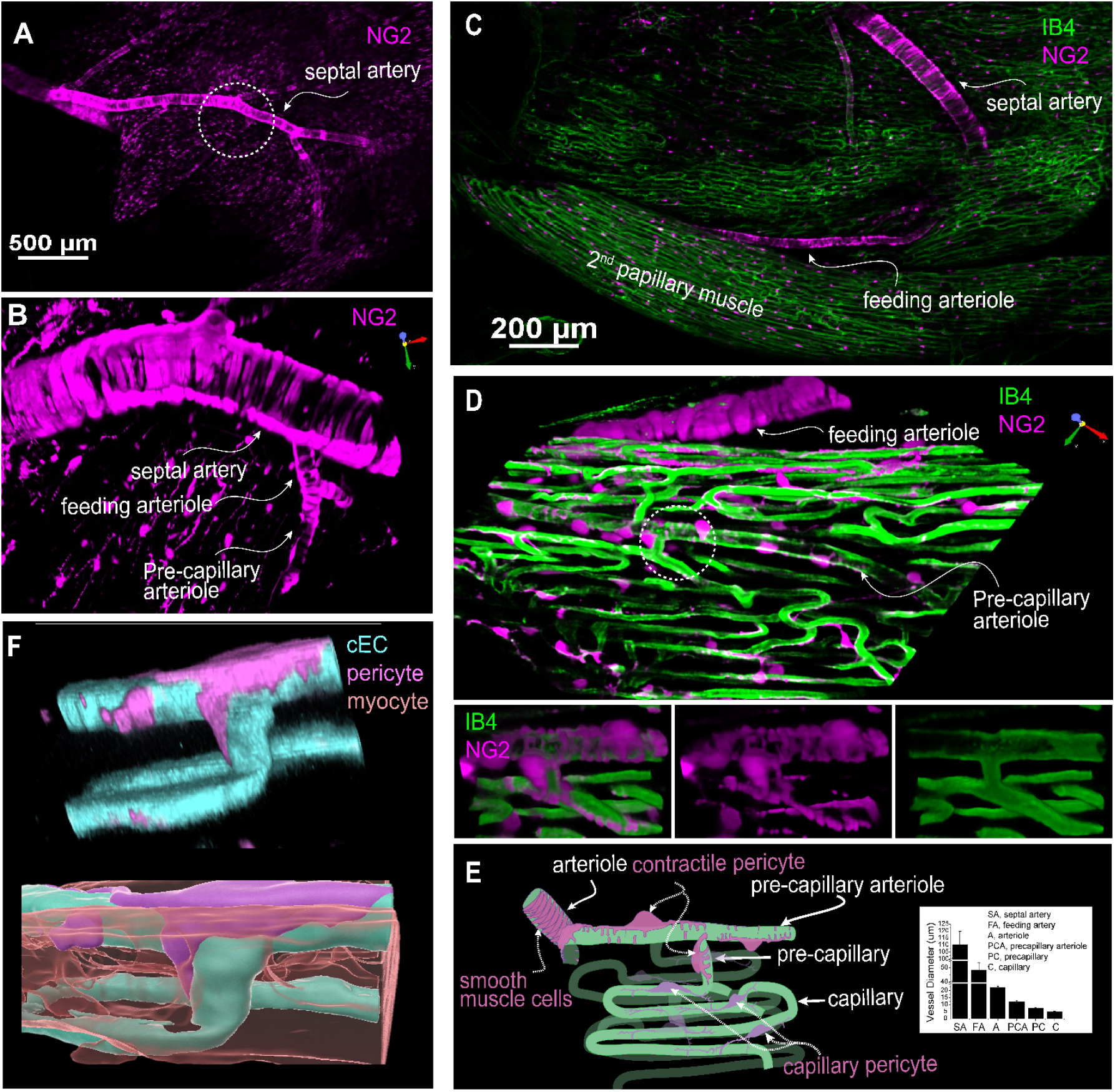
The geometry and structure of the microvascular bed of the mouse right ventricle papillary muscle. **A**. A representative image of an NG2DsRed mouse heart showing that both vascular smooth muscle cells and pericytes are NG2 positive. **B**. Zoomed-in region of the circled area in **A**, showcasing the septal artery, feeding arteriole, and precapillary arteriole through imaging with NG2DsRed. **C**. A representative image showing the branch of septal artery and feeding arteriole to the 2nd papillary muscle. **D**. Magnified 3D image of a region of the second papillary muscle showing a high-resolution image of the feeding arteriole, precapillary arterial and capillaries. Lower panel: zoomed-in image of the circled area from main panel. Note that the precapillary arteriole is sparsely covered by the contractile pericytes (often called ensheathing pericytes). **E**. A cartoon drawing that depicts the microvessels and pericytes in papillary microcirculation. Ventricular myocytes fill the “apparently empty” space between the capillaries. Inlet shows the diameter of different segment of the vessel bed at 40-80 mmHg. **F**. A pseudo-color 3D rendered image showing a pericyte, capillaries, and a myocyte (lower panel). The myocyte(s) is translucent to show its location with respect to the pericytes and cECs. **A** to **D** are from living Z-Prep in the presence of BDM; **F** is taken from a fixed heart tissue. The cEC, pericyte, and myocyte are revealed by IB4, NG2 and mitotracker, respectively.

**Figure 4.**
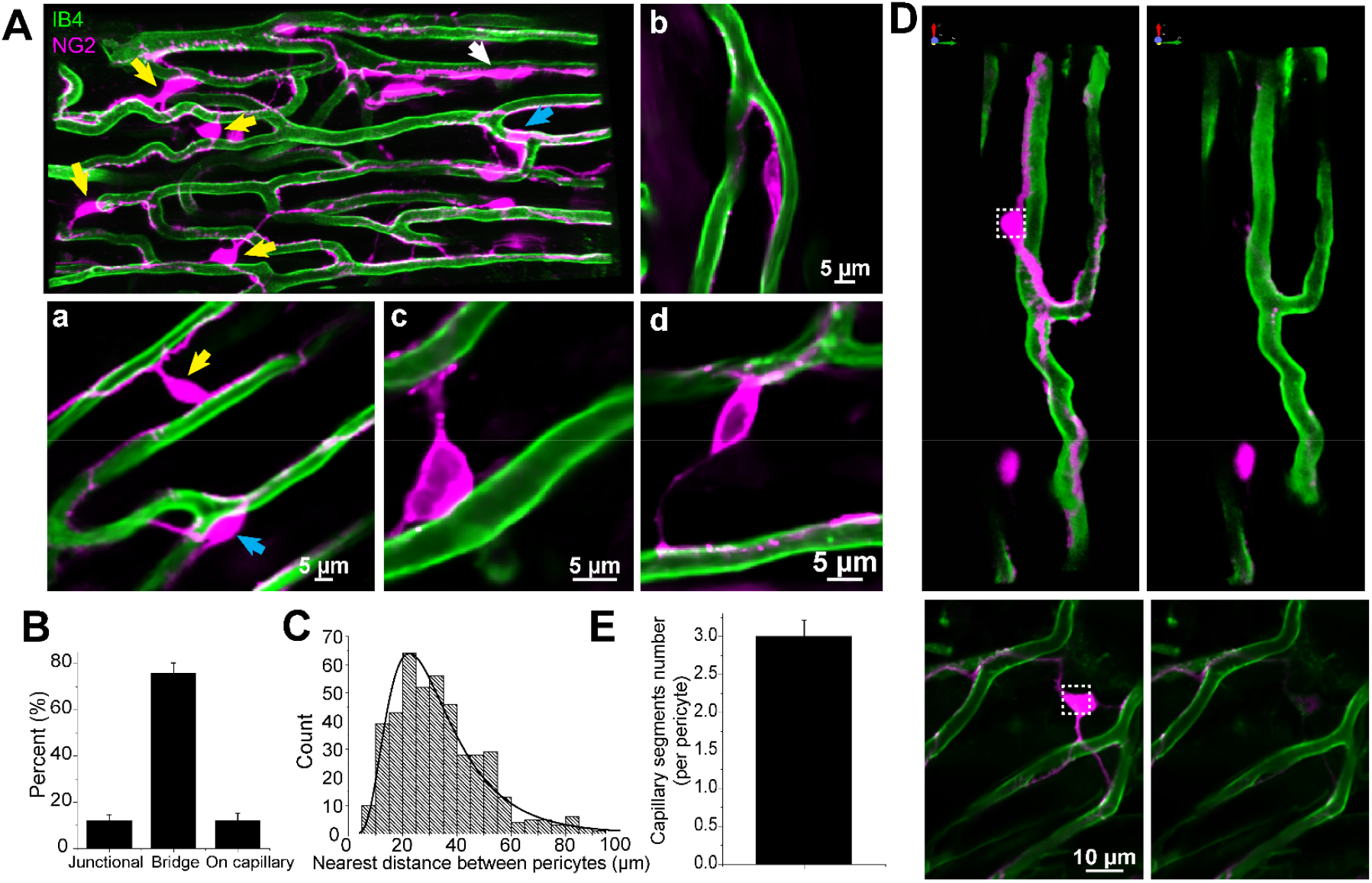
Capillary pericyte geometry and distribution in mouse right ventricle papillary muscle (Z-Prep). **A**. NG2DsRed mouse heart image displaying pericytes (magenta) and microvessels (green). Various types of pericytes are indicated by arrows: yellow for bridging pericytes located between capillaries, blue for junctional pericytes located at bifurcation area, and white for pericytes located directly on capillaries. Zoomed-in images (**A**a to **A**d) provide detailed views of pericyte localization. **A**a depicts both bridging and junctional pericytes, while Ab shows a pericyte directly attached to a capillary. **A**c and **A**d highlight bridging pericytes. Pericytes in Ac to Ad are PDGFRβ (magenta) positive cells. **B**. Summary data presenting the distribution of different pericyte subtypes. **C**. Histogram showing the nearest distance between two pericytes. **D**. Images showing pericytes and their processes before (left panels) and after photo-bleaching (right panels). **E**. Summary data showing the number of capillaries that each pericyte covers (n=10 pericytes from 2 mouse papillary muscle).

**Figure 5.**
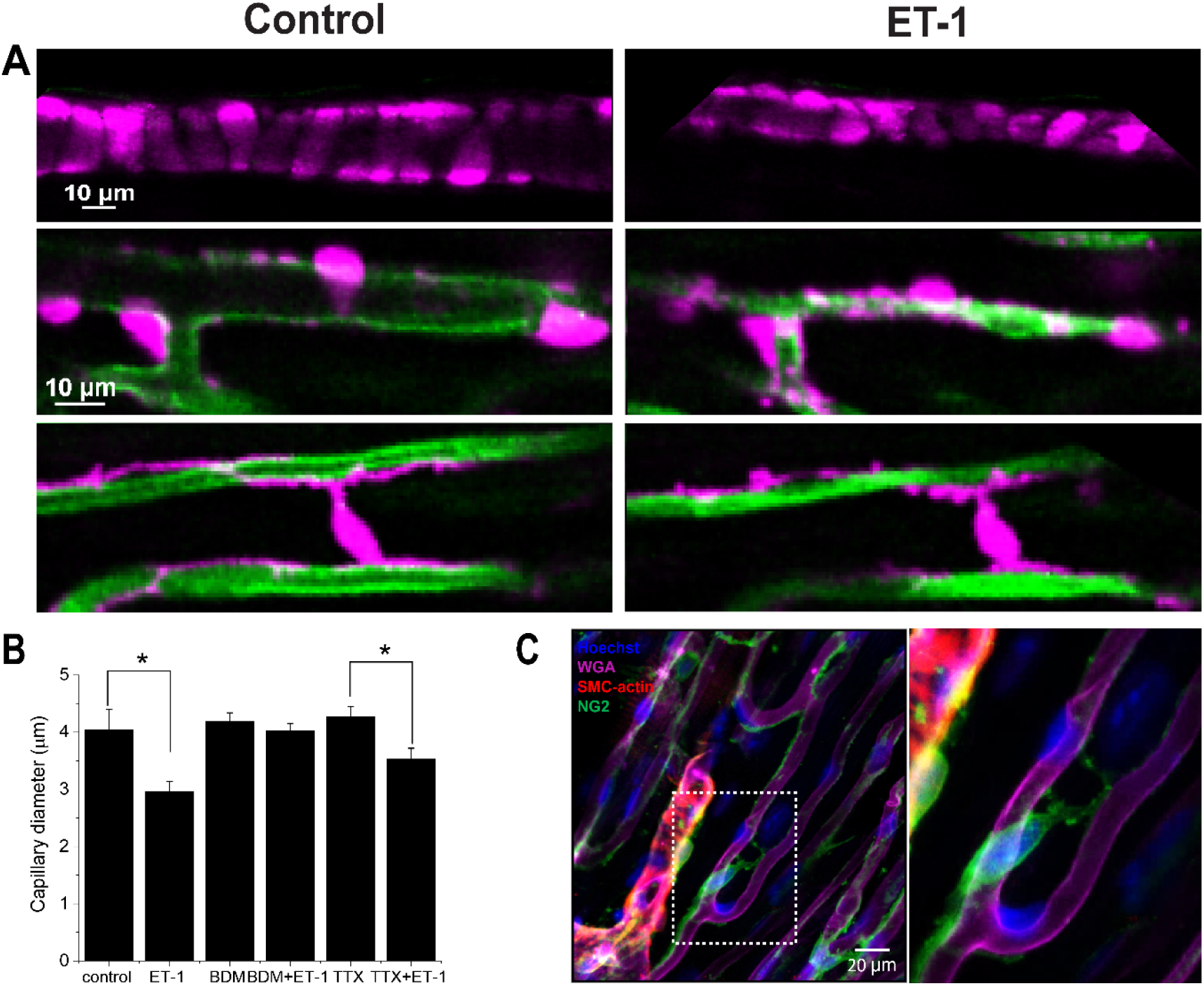
The dynamic change of cardiac pericytes and capillary. **A**. Images show a feeding arteriole (upper panel), precapillary arteriole (middle) and capillary changes in the presence (left) and absence (right) of BDM on top of ET-1. These images were obtained from the same Z-Prep as in **Figure 3C. B**. Summary data show capillary diameter change in the presence of ET-1 and/or TTX. *p<0.05. **C**. Whole-mount immunostaining reveals that smooth muscle actin (SMC-actin, red) is expressed in pre-capillary arteriolar smooth muscle cells but not in pericytes (green). The papillary muscle was stained using anti-NG2 (green), anti-SMC-actin (red), WGA (magenta), and Hoechst to identify pericytes, smooth muscle cells, endothelial cells, and nuclei, respectively. Right panel, zoomed-in image of the boxed area in the left panel.

**Figure 6.**
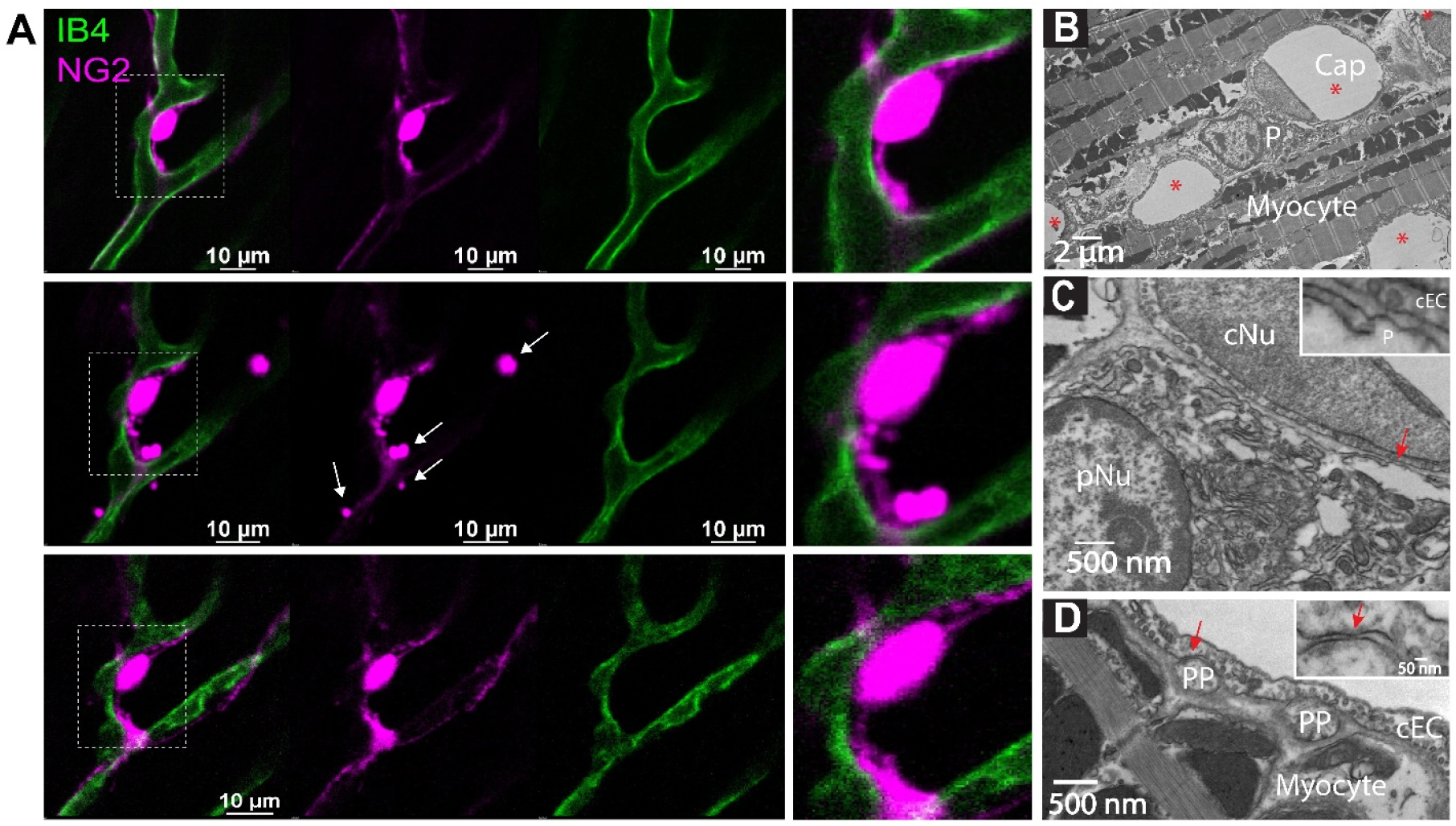
Pericytes Are Not Necessarily Involved in Capillary No-Reflow. **A**. Images showing the change and restoration of pericyte structure in the presence and absence of ET-1. Note the granulation of pericyte processes (magenta, white arrows) in the presence of ET-1 (middle panel) and the restoration after washing out ET-1 (lower panel), while the capillary lumen remains unaffected. The right panels provide a zoomed-in view of the boxed areas in the left panels. **B-D**. Transmission Electron Microscopy (TEM) myograph illustrating the structural interaction between pericytes and capillary endothelial cells (cEC) in the mouse heart. **B**. Cross section of mouse papillary muscle showing five capillaries (Cap, *) and one pericyte soma (P). cNu, capillary endothelial cell nucleus; pNu, pericyte nucleus. Cap, capillary. **C**. Magnification of the framed area in **B**. The inset zooms in on the arrowed area in **C**, showing the distance between the pericyte soma membrane and the cEC membrane. **D**. Cross section highlighting two pericyte processes (PP). The inset magnifies the junctional structure between the pericyte process membrane and the cEC membrane.

### The surprisingly high abundance of pericytes

We also measured the volume and the number of pericytes relative to capillary segment length and the number of cardiac myocytes. The total volume of the pericytes including processes occupies ∼4.8 ±0.06 % of the total tissue volume, or 1/3 of the total capillary volume. The quantification of pericytes was conducted based on the total volume of the imaged muscle. On average, there are approximately 15 pericytes per 10^6^ μm^3^ of papillary tissue, indicating one pericyte for every two to three cardiac myocytes! This calculation considers the mean volume of a single myocyte, which measures about 28,894 ± 3,875 μm^3^. Here there is an estimated density of approximately 35 myocytes per 10^6^ μm^3^ of papillary muscle tissue.

### Location and “reach” of pericytes

Additionally, we determined the pericyte abundance along the capillary segments. On average, there is one pericyte every ∼150 μm along mouse capillaries. (This differs from previous reports on rat left ventricles which showed a greater abundance -- one pericyte every 60 μm ^10^.) Given that the average capillary segment length in mouse is 58.7 ± 3.8 μm (**Figure 2**), it follows that each pericyte and its processes could reach up to three capillary segments but this “reach” depends on the local geometry. Note that this estimated “reach” of a pericyte was confirmed through photo bleaching experiments (**Figure 4D** and **4E**). Moreover, we observed that the nearest distance between two pericyte somas is on average 32.5 ± 1.4 μm (n=115 pairs of pericytes), which is greater than the nearest distance between inter-capillary distance ^9, 26^.

### Pericyte contraction and microvascular blood flow regulation

That pericytes contract and thereby narrow the diameter of capillaries to regulate blood flow is established in diverse tissues including brain ^10, 35-37^. To examine the responsiveness of capillaries and the mechanisms under which the capillary diameters change, we applied endothelin-1 (ET-1) (e.g. 5-10 nM) in the vascular lumen. This application led to significant constriction in capillaries and other vascular elements. In our Z-prep experiments it is important to minimize motion of the Z-prep. In many experiments we have used BDM to reduce or block movement produced by contracting ventricular myocytes. In the experiments shown here using ET-1 to produce contractions of pericytes, we needed to use another method to block movement of ventricular myocytes because BDM also blocks the action of ET-1 on pericytes. To block contractions of ventricular myocytes without blocking the action of ET-1 on pericytes, we used TTX (300 nM to 1 uM). TTX is an ultra-specific inhibitor of Na^+^ channels ^38^ and effectively stops action potential (AP) induced contractions of ventricular myocytes ^39^ (**Figure 5B**). TTX thus stops cardiac contractions without affecting pericytes or capillary endothelial cells, cell types that have no voltage-gated Na^+^ channels. [Note: Even if there were voltage-gated Na^+^ channels in cEC, pericytes or smooth muscle cells, these Na^+^ channels would be inactivated due to the relatively positive membrane potential of these cells (∼-35 mV).] Therefore, the application of TTX should not significantly affect vascular function under physiological conditions ^40-42^. As shown in the **Figure 5 and Figure 6**, ET-1 (5-10 nM) caused both arterioles and capillaries to constrict under these conditions. Interestingly, for the precapillary arteriole, the constriction is likely attributable to the processes (**Figure 5A**, upper panel), whose constriction almost closes the arteriole. For the capillaries, ET-1 induces the cell body of bridging pericytes to shorten and round-up (**Figure 5A**, lower panel). In the presence of ET-1 the processes of all the pericytes developed a “granulated” or “lumpy-bumpy” appearance (**Figure 5** and **Figure 6**). The mechanisms that underlie this morphological transformation and the contraction remain unknown, as no alpha smooth muscle actin was detected in capillary pericytes (**Figure 5C**), consistent with the observations in other tissues including brain^34, 37, 43^.

### Intercellular connections in heart

Enhanced specific communications between pericytes and capillary endothelial cells are reported in the brain at macroscopic sites called “peg and socket” connections. However, the exact nature and abundance of the enhanced communication is suggested by the presence of gap junctions in pericytes and endothelial cells and rare couplings consistent with clusters of gap junctions ^44^. There does not appear to be evidence of specific electrical synapses or of one or a cluster of gap junctions. For us to investigate the physical and signaling relationship between cEC’s and pericytes in heart, we used the tools of transmission electron microscopy. This was done using the same mouse right ventricle papillary muscle preparation as the Z-prep. **Figures 6B to 6D** show our findings from this investigation. First we did not find evidence of “peg-and-socket” connections in contrast to reports for brain microvessels ^21, 45-47^. There was evidence that the neighboring cells (pericytes and cEC’s) had “intimate” regions where both cells were as close to each other as was seen in peg-and-socket connections. Among the 67 pericyte cross sections (including processes or soma) in 19 capillary segments, 14 displayed regions that could have contained gap junctional structures (**Figure 6C** and **6D**), where the closest distance between the capillary endothelial cell (cEC) membrane and the membrane of pericyte processes was less than 20 nm. However, the closest distance between the pericyte soma and the cEC membrane was approximately 40 nm (**Figure 6C** and **6D**), a distance too large to support a gap junctional structure. So, like the data from brain and related peg-and-socket systems, perhaps it is the pericyte processes where pericytes communicate with cEC’s. To state this clearly, the mechanisms by which pericytes regulate capillary diameter and microcirculation remains unclear, despite the apparent roles of pericytes in ischemic “no-reflow” in the heart ^10^. As depicted in **Figure 5C** and supported by previous research, capillary pericytes do not express measurable smooth muscle actin, the primary filament protein responsible for constriction in smooth muscle cells and contractile pericytes ^34, 37, 43^. However, the application of ET-1 did induce a decrease in capillary diameter as well as the morphological change of the pericyte processes, as shown in **Figure 5** and **6**. Thus, we have reason to believe that the decrease of capillary diameter is due to the structural and/or functional change of pericytes. However, surprisingly, after washing out ET-1, we found that the constriction of capillary remains, while the structure of pericytes and their processes seemed restored (**Figure 6A**). These data might suggest that 1) capillary endothelial cells were damaged permanently; or 2) the change of pericyte morphology and capillary diameter are two independent “ischemic reactions”. As shown in **Figure 5A**, the precapillary arterioles constrict completely which might result in the “stopping of flow” in the downstream capillaries ^48^.

## Discussion

Pericytes are small pleomorphic cells associated primarily with capillaries and appear to have a distinct set of roles in regulating blood flow in heart and in other tissue^4, 30, 34, 37, 49, 50^. They are attached to the outside of the capillary endothelial cells that form the capillaries -- the small tube-like structures of the smallest diameter element of the vascular system. This location places them on the exterior of the capillary wall which makes them “mural” cells analogous to the vascular smooth muscle cells that surround the somewhat larger arterioles and end-arterioles that connect the blood flow supply to the capillaries themselves. Published evidence identifies multiple morphologically distinct pericytes types in the brain that are associated with specific branches of the capillary tree ^24, 30, 34, 51, 52^. Published findings suggest electrical connections between pericytes and cEC’s through a structural entity called a “peg and socket” junction ^44, 53^. However, the molecular connection between the different cell-types remains to be established. Both cell-types have a membrane potential of about -35 mV ^2, 54^. Here we have presented images of cardiac pericytes that look like pericytes in other tissues with respect to their organization on the cEC’s. There are two wide-spread similarities. A typical pericyte frequently shows a “bump on a log” nuclear protrusion with cytoplasmic thin strands that travel along part of the length of the capillary (tens of microns) ^11, 55^ (**Figure 3**). In brain and heart there are NG2 positive pericyte structures that may sit at the bifurcation of a small arteriole that splits into two capillaries ^4, 34^. These structures appear to function as a directionally controlling pre-capillary sphincter ^34^. However, there are also some important differences. Specifically, the cardiac pericytes send extensions directly to nearby capillaries and also to more distant capillaries with these longer extensions traveling on the surface of ventricular myocytes or between them. We have also identified the appearance of pre-capillary “sphincters” in heart that appear to be a transition cell type resembling both single smooth muscle cells and also pericytes. They may entwine bifurcating arterioles as they transition to two outbound capillaries (**Figure 3D**). While there is evidence that the capillary pericytes can contract ^37, 56^, there is insufficient data at the present time to characterize the full nature of the control of this contraction and their kinetics in heart.

The imaging of pericytes in heart shown here raises several challenging questions that some are discussed below. First, how does the observed distinctive anatomy of the cardiac pericytes suggest they work in the hypothesized electro-metabolic signaling of heart? ^2-4^. What experiments could eventually be done to test these ideas? What can we learn about pericytes from their contractile behavior following the application of endothelin in heart and brain? Does the concept of “no re-flow” provide us with understanding of pericyte function and how does this relate to gross changes in pericyte morphology? What does the diversity of pericyte extension length and shape suggest as pericyte extensions traverse heart tissue?

### Electro-metabolic signaling (EMS)

Investigating how the metabolic states of heart muscle are linked to the regulation of blood flow is a huge experimental challenge. Our current understanding of the EMS hypothesis has been recently reviewed ^3^. Active investigations involve measuring electrical and chemical messages found in vascular myocytes, capillary endothelial cells, pericytes as well as those from ventricular myocytes. How these interactions contribute to cardiac physiology and pathology has the potential to reveal the links between changes in the metabolic activity of the myocytes and the control of blood flow. Such measurements involve local and regional signals but, to the extent the data are accurate, they should broaden our understanding of cardiovascular physiology and pathology. However, this effort demands innovative techniques and other methods that are still under development. Considering these major challenges, this present study was conducted to provide more precise information on the geometric arrangement and dimensions of myocardial precapillary arterioles, capillaries and venules, as well as the architecture, function and categories of pericytes in the heart. Such work will enable our use of more accurate values and terms in models that describe the microcirculation in the heart. Through careful experimentation using innovative methods, we described the structure of microcirculation, the diversity of pericytes, and their functional implications in mouse papillary muscle. Collectively, our new results contribute to a more comprehensive understanding of the organizational hierarchy of capillaries, the structural diversity of cardiac pericytes, their strategic localization, and their functional roles within the heart. Thanks to the living work model, the Z-prep, additional findings have been made. While much of the new data agree with the previous work, there are surprises. This research thus is an extension of our previous work ^1-4^, providing structural foundation for the electro-metabolic signaling regulation of blood flow in heart.

### Microcirculatory structure in heart

Following arteries and arterioles, where the pattern of smooth muscle cells is clear, fewer smooth muscle cells are seen in the branching vessels. Instead, the vessels are covered with pericytes (**Figure 3**). While both smooth muscle cells and pericytes are NG2-positive, smooth muscle cells are short, ring-shaped and densely packed, which are easily identified and distinguished from pericytes, especially in situ (**Figure 3 and Figure 4**). Some researchers refer to all the vessels that are covered by pericytes as capillaries, because pericytes were initially defined as cells that decorate capillaries ^23, 55, 57-59^. Others call the vessels with “ensheathing” pericyte proximal capillary ^30, 51^ or pre-capillary arteriole because these pericytes are smooth muscle actin-positive and function similarly to smooth muscle cells ^24, 34, 52^. Our observations support the latter concept and called the vessels covered with contractile pericytes pre-capillary arteriole (**Figure 3**). Accordingly, the pericytes on the pre-capillaries are referred as “contractile pericytes” or “pre-capillary pericytes”. A specific single contractile pericyte that is located on a short pre-capillary segment is thereby referred as pre-capillary “sphincter”. Unlike what has been observed in the brain and other tissues where capillary arrangements are cascaded in a sequence, in the heart, capillaries are distributed relatively regularly and at a single “level” (**Figure 2** to **4**) in “working” myocardium. These capillaries are primarily oriented parallel to the long axis of the cardiac myocytes and are inter-connected with short perpendicular stretches (see **Figure 2**), similar to the arrangement found in skeletal muscle ^60-62^. Thus small diameter capillaries run parallel to the myocytes with branches at approximately right angles. We do not recommend renaming the cardiac capillaries with order numbers as done by Kassab and his group ^12, 13, 63^. Re-stating the organization -- there is primarily a single capillary type serving functioning heart muscle but with two orientations. In brain, the complexity is much greater. The cardiac capillaries have been morphologically divided into subtypes based on their three locations: pre- and post-capillary and working capillary with appropriate pericytes for each type ^11^. Our work shows that there are actually two main functional types of capillaries for the functioning/working myocardium, one runs parallel to the myocytes and is mainly on the capillary and another called a “bridging pericyte” that starts on one capillary and passes intimately by a myocyte and ends on a parallel capillary. All the capillaries have similar diameter regardless of the position or segment length.

### Cardiac pericytes morphology, classification and blood flow regulation

While the contractile properties of pericytes have been studied quite thoroughly in other tissues, such as in the brain, research on the fundamental functions of pericytes in the heart is still lacking. Most studies have focused on changes in pericytes under pathological conditions such as heart failure and myocardial ischemia. One reason for this is the constant beating of the heart, which makes it challenging to observe real-time changes in pericytes— precisely what is needed to study pericyte contractility. Thanks to our new research model, we can now monitor the dynamic changes of the capillary diameter and pericyte morphology. Our imaging of live tissue clearly shows the variation in the capillary diameter and the relationship between the diameter and the surrounding pericytes. We found that local narrowing of capillaries occurs more often in the absence of a vasodilator. In addition, the narrow areas do not necessarily coincide with the location of pericyte somas, which differs from the findings reported by Attwell’s group and others ^10^. They found that ischemia reperfusion caused pericytes to contract and resulted in no-reflow in heart. However, their data come from tissue that is not actively perfused. A large number of studies suggested that pericyte contraction reduces capillary flow and decreases capillary diameter. On the other hand, damage or loss of pericytes may lead to the stop (or “no reflow”) or insufficiency of blood flow ^10, 29, 49, 64, 65^. Our observation in situ heart tissue strongly suggest that the constriction of capillary is secondary to the constriction of their upstream arterioles, which control the downstream blood flow. When feeding with the perfusates including ET-1, arterioles are the first to response and constrict. This is consistent with the report in the brain ^52^. Later studies demonstrate that longer and stronger depolarizing stimulation would enable capillary pericyte to contract ^52^, suggesting that capillary pericytes are not necessarily involved in capillary blood flow regulation under physiological conditions ^66^. The diameter change starts a while after the arteriole constricted, suggesting that either the arteriolar smooth muscle or precapillary pericytes are more sensitive to ET-1 or other vasoconstrictors. Indeed, the circular processes are rich with contractile filaments (**Figure 5**) ^34, 51^. Channelrhodopsin excitation contracts the brain pericytes and reduce the blood flow ^67^. At present, the literatures are sort of in agreement that all the pericytes are contractile regardless of their location, although the pericytes along the brain capillary contract at a much slower rate, exerting a more gradual effect on blood flow regulation ^37^. However, only very few observations made on capillary pericyte (thin strand pericyte) support this conclusion. Most of the measurements were conducted on the “capillaries” with ensheathing or mesh pericytes which are smooth muscle actin positive ^28, 34, 49^. The diameter of these “capillaries” is generally > 6 μm ^28, 29, 59, 68^, which fall into our precapillary arteriole category ^24, 49^. We therefore refer to the pericytes on the pre-capillary arteriole as “contractile pericytes” to acknowledge their properties, functions, and location. This variety in pericyte form and placement reflects a sophisticated level of vascular regulation, hinting at a complex system where each pericyte subtype may be finely tuned to meet the specific demands of the tissue it serves.

### Pre-capillary sphincters, smooth muscle cells and pericytes

Originating in 1937 when Zweifach and his colleagues described the precapillary sphincter in the mesentery of frogs ^69^, this particular structure has long been thought to be universally present in the microvessels of all tissues and has been included in physiology textbooks. However, this structure is typically depicted through artistic drawings in textbooks rather than from original experimental observations. Traditionally described as a ring of smooth muscle at the arteriolar-capillary junction, sphincters are indicated to regulate blood flow into the capillaries. Even though the structure and function of the precapillary sphincter were hot topics and had been widely studied in the last century ^68, 70-73^, the precapillary sphincter was not well-defined due to technical limitations. Firstly, three-dimensional imaging was unavailable, making it difficult to accurately illustrate the sphincter’s morphology. Secondly, vessels in *ex vivo* or *in vitro* preparations were not perfused, so their diameters did not reflect physiological conditions. Lastly, there were no biomarkers to define the cell types. Although these technical limitations have now been resolved, the definition, territory, structure, and function of the precapillary sphincters remain unclear, especially in the heart. Recent research indicates that the precapillary sphincter in brain regulates local blood flow ^58^, and its loss or impaired dynamics might cause neurovascular dysfunction^58 57, 58, 74, 75^. Among all smooth muscle cells or pericytes capable of constriction, it exhibits the greatest contraction upon stimulation ^58^. In skeletal muscle, where the demand for oxygen can fluctuate dramatically, these sphincters are also postulated to play a critical role in directing blood flow in response to local metabolic needs. However, the existence and functional role of precapillary sphincters in the heart have never been clear. In fact the existence and the function of sphincter has been a topic of debate among vascular biologists for many years ^76^. Some researchers even question whether this structure is indeed present in tissues other than the mesentery and have even pointed out that it is not universally present ^76, 77^. In this study, we have identified sphincter-like structures in the heart, as shown in **Figures 3**. These structures consist of one or a few pericytes located between capillaries and arterioles. While some scholars refer these pericytes as “mesh pericytes” ^4, 23, 24, 30^, we prefer to name them “contractile pericytes” or “precapillary pericytes” ^4^. They are among the last group of pericytes that express smooth muscle actin with high contractile activity. Even though it is still unclear how this type of pericyte regulates blood flow into the downstream capillaries, the ET-1-induced constriction (**Figure 4**) of this pericyte was observed and the presence of α-smooth muscle actin was confirmed ^4^. Given that the classical concept of the precapillary sphincter may not apply uniformly across all tissues, the fundamental principle of blood flow regulation at the microvascular junction is a critical aspect of tissue physiology that may be carried out by various mechanisms across different organ systems.

In summary, using perfused living cardiac muscle, this study has described the geometry and fine structure of the microvasculature and its components in the mouse heart, with an emphasis on the organization and diversity of pericytes. Three pericyte identities appear to exist in heart: pre-capillary sphincter pericytes, longitudinal pericytes and bridging pericytes. This report thus lays the foundation for future functional studies and provides guidance for identifying mural cell types on and around the capillaries in heart. Additionally, our real-time imaging enables dynamic tracking of capillary diameter changes under stress and challenges in heart. Finally, this work provides the organizational foundation for future studies on the physiological roles of cardiac pericytes and some basic guidance on how they may work.

